# Epistasis at the cell surface: what is the role of Erg3 loss-of-function in acquired echinocandin resistance?

**DOI:** 10.1101/2025.05.08.652905

**Authors:** Hans Carolus, Judith Díaz-García, Vladislav Biriukov, Stef Jacobs, Dimitrios Sofras, Alicia Pageau, Celia Lobo Romero, Lore Vinken, Pilar Escribano, Jesús Guinea, Katrien Lagrou, Christian R. Landry, Toni Gabaldón, Patrick Van Dijck

## Abstract

Echinocandins, which target the fungal β-1,3-glucan synthase (Fks), are essential for treating invasive fungal infections, yet resistance is increasingly reported. While resistance typically arises through mutations in Fks hotspots, emerging evidence suggests a contributing role of changes in membrane sterol composition due to *ERG3* mutations. Here, we present a clinical case of *Nakaseomyces glabratus* (*Candida glabrata*) in which combined mutations in *ERG3* and *FKS2*, but not *FKS2* alone, appear to confer echinocandin resistance. Integrated analyses reveal a recurrent association between Erg3 loss-of-function and echinocandin resistance mediated by Fks variation across *Candida* species, but exclude *ERG3* loss-of-function as an independent resistance mechanism. Advances in Fks structural biology and insights into echinocandin-Fks interactions support a model of epistatic crosstalk between membrane sterols and Fks function. Understanding this interaction is crucial, as it may underlie not only acquired echinocandin resistance but also the broader development of multidrug resistance across major antifungal drug classes.

## Main

Echinocandins are a first⍰line drug class for treating invasive fungal infections. They inhibit the fungal-specific enzyme β-1,3-glucan synthase (Fks), disrupting glucan synthesis and cell wall integrity. Resistance to echinocandins typically arises from mutations in *FKS* genes, mainly accumulating within three specific mutational hotspot (HS) regions of Fks (1, 2). However, recent evidence suggests that sterol-mediated alterations in membrane composition can also modulate echinocandin susceptibility. This study presents a clinical case of *Nakaseomyces glabratus* supporting this concept. Through strain analyses, data mining, and literature review, we explore the hypothesis that the loss-of-function (LoF) of a C-5 sterol desaturase (Erg3), which alters the membrane sterol composition, influences echinocandin resistance in the context of Fks modulation through allostery. Understanding the putative epistatic interactions between *ERG3* and *FKS* mutations is essential, as it may significantly drive the evolution of multidrug resistance and challenge therapeutic efficacy.

### Erg3-Fks cross-talk: an intruiging clinical case of *N. glabratus*

We re-analyzed two clinical isolates of *N. glabratus* from a peritoneal abscess (isolate A) and the bloodstream (isolate B) of a patient admitted to the Hospital General Universitario Gregorio Marañón (Madrid, Spain) in 2020 (3). Both *in vitro* and *in vivo* susceptibility testing demonstrate that isolate B but not isolate A was resistant to echinocandins (**Figure 1**). There was a 5.5-fold and 7.9-fold increase in micafungin and anidulafungin MIC values, in isolate B compared to isolate A, respectively (**Figure 1A**). Additionally, isolate A showed significantly reduced *in vivo* colonization under micafugin treatment in all organs in a murine infection model, while isolate B seemed insensitive to micafungin treatment (**Figure 1B**).

**Figure 1.**
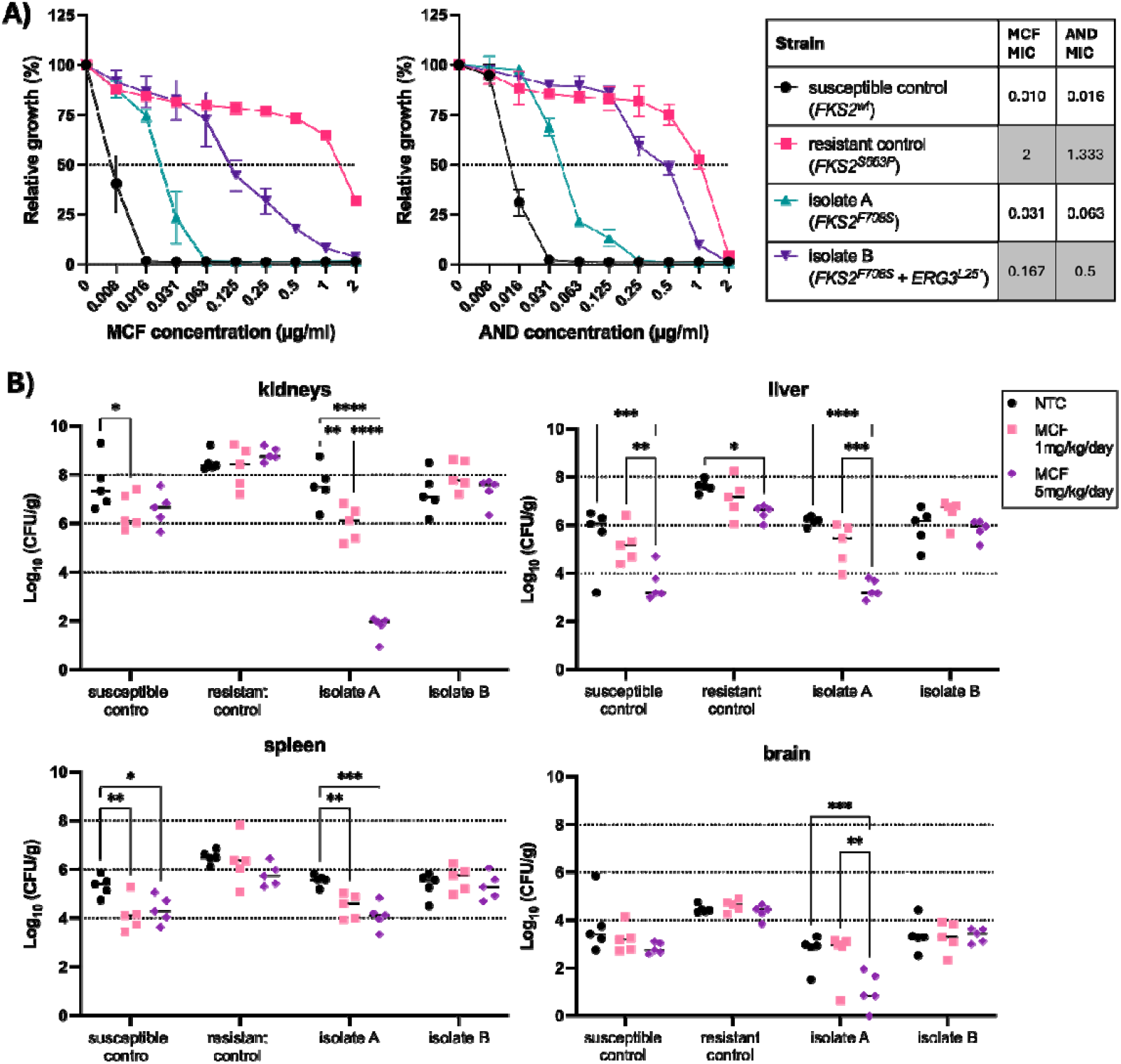
*In vitro* and *in vivo* echinocandin susceptibility of the reported clinical strains (isolate A and B) and a susceptible and resistant control strain of *N. glabratus*. Strain details are described in the *Methods* section. **A)** Broth dilution susceptibility test for micafungin (MCF) and anidulafungin (AND) was performed according to the EUCAST method (4) on three independent subcultures per isolate. The table shows the mean value of minimum inhibitory concentration (MIC) of 50% growth, with grey indicating above-breakpoint resistance according to EUCAST clinical breakpoints (5). **B)** *In vivo* echinocandin susceptibility evaluation was conducted in an immunocompromised murine systemic infection model. Mice were treated with 2 doses of MCF (1 mg/kg/day and 5 mg/kg/day) or a PBS vehicle (NTC: non-treated control) for 7 days, after which organ colonization was assessed by CFU enumeration from organ homogenate plating. Significant differences (two-way ANOVA, Tukey’s test) of pairwise comparisons are shown, with *:P ≤ 0.05; **:P ≤ 0.01; ***: P ≤ 0.001 and ****:P ≤ 0.0001.

A whole genome sequence comparison of isolates A and B revealed 42 unique nonsynonymous mutations across 29 genes (**Supplementary Table S1**). Both strains harbored the same mutation (F708S) in *FKS2*, outside of HS1, while isolate B additionally contained a putative LoF mutation in *ERG3* (L25*). Based on a literature review, no additional genetic variants besides those in *FKS2* and *ERG3* could be associated with echinocandin resistance. These findings suggest that the putative LoF of Erg3 might contribute to echinocandin resistance in the *FKS2*-mutated background of isolate B.

### LoF mutations in *ERG3* are commonly associated with echinocandin resistance

Next, we further explored how common combined *ERG3* and *FKS* variation is in the context of echinocandin resistance by mining the recently constructed fungAMR database, which curates literature-reported mutations associated with antifungal drug resistance (6). We found reports of the combination of *FKS1/2* and *ERG3* variation in multiple *Candida* species, including *Candidozyma auris* (7, 8), *Candida albicans* (9, 10), *Candida lusitaniae* (11) and *N. glabratus* (12, 13), often isolated after exposure to echinocandins only (**Table 1**). All 31 *ERG3-FKS* variants were reported to be resistant to at least one echinocandin. Moreover, in serial clinical isolates of *N. glabratus* investigated by Lim *et al*. (12), the combination of *ERG3*^*G236D*^, *ERG3*^*W98\**,^ and *ERG3*^*F226X*^ mutations with *FKS2*^*L1357E*^ and *FKS2*^*FL659L*^ mutations reduced the susceptibility to echinocandins, compared to strains with only *FKS2* mutations. For instance, the MIC values for caspofungin, micafungin, and anidulafungin increased over 32-fold in an isolate with both *FKS2*^*FL659L*^ and *ERG3*^*W98\**^ mutations, compared to an isolate with an *FKS2*^*FL659L*^ mutation alone. Furthermore, the combination of an *FKS2*^*L1357E*^ and *ERG3*^*F226X*^ mutation conferred anidulafungin resistance, which was not present in an isolate of the same clinical background, with an *FKS2*^*L1357E*^ mutation alone (12). Similarly, the acquisition of an *ERG3*^*L207I*^ mutation further reduced the susceptibility to caspofungin, in an already resistant *FKS1*^*FL635L*^ *&* ^*M690I*^ mutant of *C. auris* (8). Ksiezopolska *et al*. (13) found the co-occurrence of mutations in *ERG3* and *FKS1* and/or *FKS2* in 19 *N. glabratus* strains, representing 25% of the total number of sequenced strains that were experimentally evolved in anidulafungin.

**Table 1.**
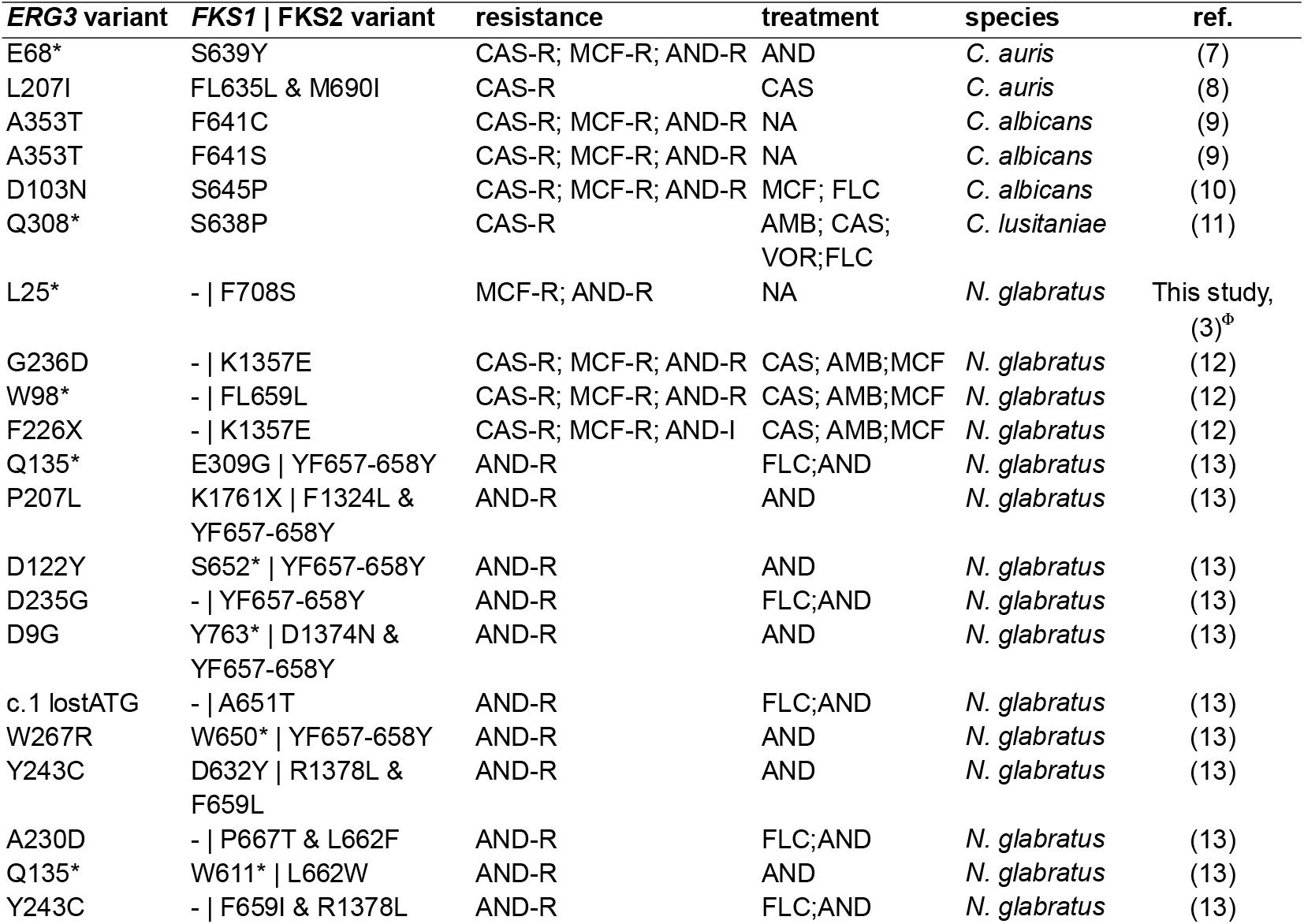
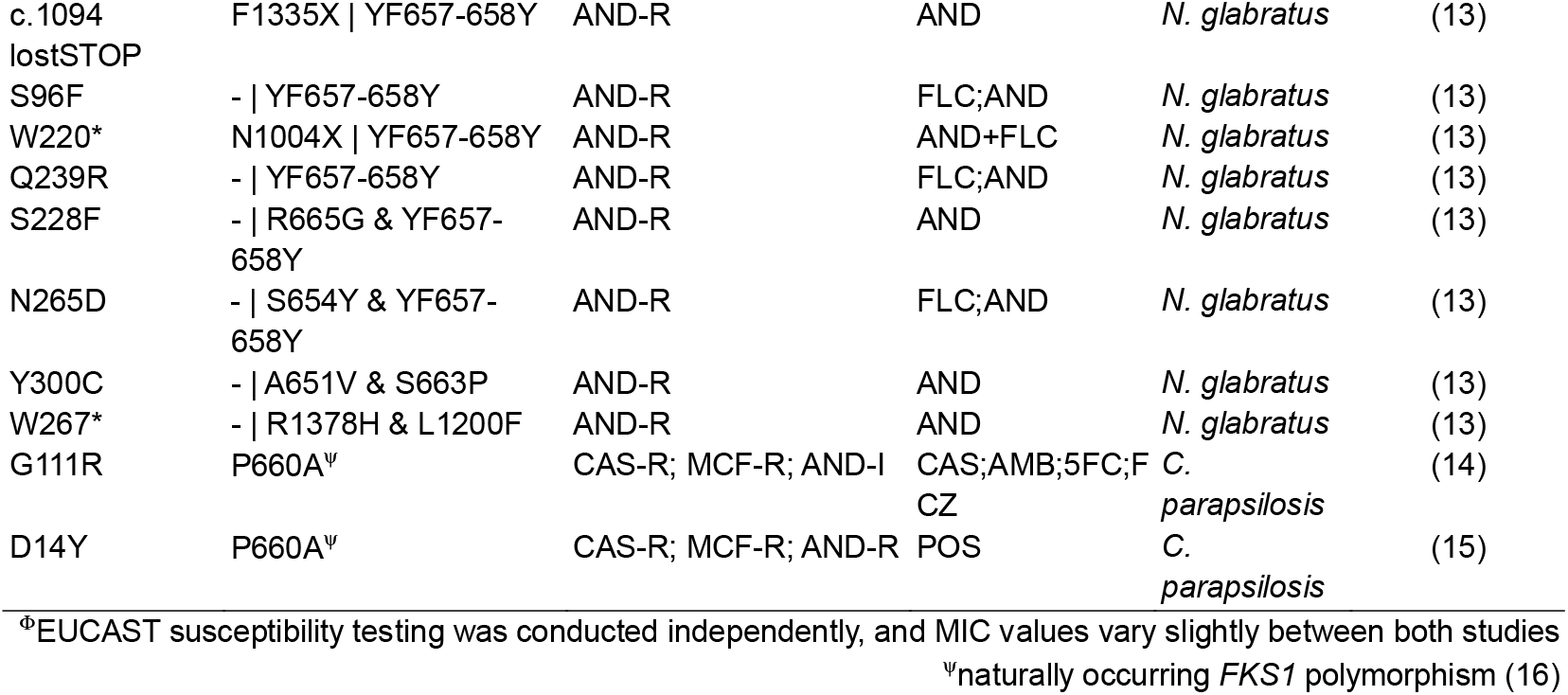
Combined variation in *ERG3* and *FKS* orthologues across *Candida* species, as reported in FungAMR (6). Mutations are noted as the amino acid changes in the species as reported in the corresponding references. Only reports that include both *FKS1/2* and *ERG3* genotyping, together with echinocandin susceptibility information, were considered. The resistant (−R), or intermediate (−I) phenotype to all echinocandins assessed in the corresponding studies, is reported. CAS: caspofungin; MCF: micafungin; AND: anidulafungin; FLC: fluconazole; AMB: amphotericin B; VOR: voriconazole; 5FC: flucytosine; POS: posaconazole. NA: not reported.

Out of 31 *ERG3* variants detected in *FKS*-mutated backgrounds, 28 were unique and 10 (35.7%) were nonsense, frameshift, start-codon-loss, or stop-codon-loss mutations, implying loss of Erg3 function (**Table 1**). To assess the impact of all reported *ERG3* variants, including missense variants, we calculated an Evolutionary Scale Modeling (EMS) impact score following Brandes *et al*. (17). ESM uses an unsupervised deep-learning model trained on more than 250 million protein sequences to capture evolutionary and structural features directly from sequence data (30). **Figure 2** shows ESM impact scores for all 28 unique mutations and compares them to the EMS impact score of all *ERG3* mutations (n = 37 711) curated in the fungAMR database (6). A two-component Gaussian mixture model defines a threshold for putative LoF variants (17) at ESM score -7.88. 22 out of 28 *ERG3* mutations (78.6%) show an EMS score below this threshold, indicating that most of the *ERG3* mutations co-occurring with *FKS* mutations likely cause a LoF of Erg3.

**Figure 2.**
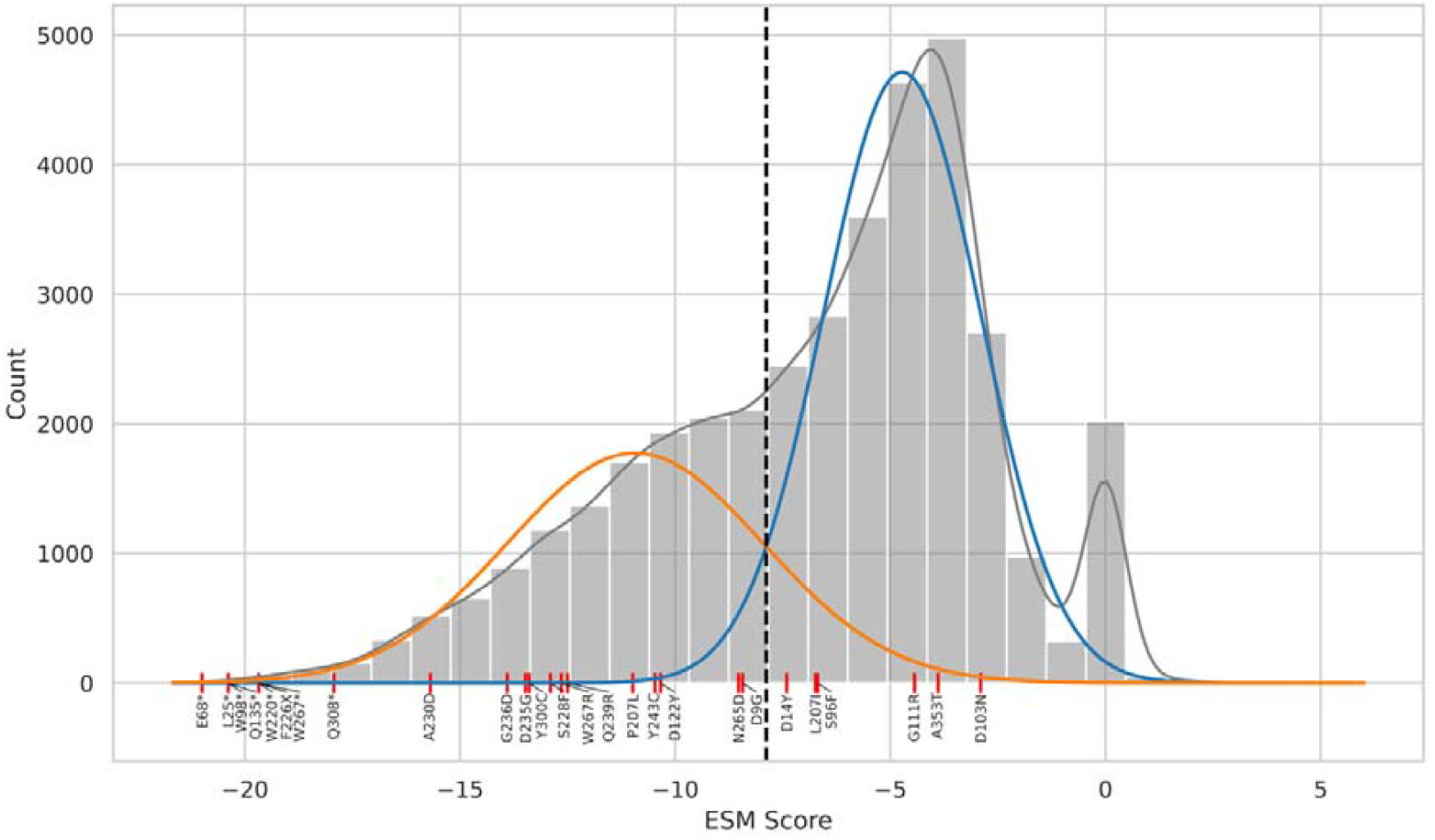
Distribution of ESM scores across 37 711 *ERG3* mutations curated in fungAMR (6) (grey), compared to variants listed in **Table 1** (red lines). 26 of 28 variants are shown, as no ESM could be calculated for the start and stop codon loss mutations. Lower ESM scores indicate a higher likelihood of the variant being damaging to protein function, as exemplified by nonsense mutations appearing at the lower extreme of the distribution. Synonymous mutations cluster around a score of 0. A two-component Gaussian mixture model was fit on the distribution, excluding synonymous mutations (blue and orange lines) (17). The intersection of both fits is -7.88 (dotted line), defining a threshold for variants that are likely LOF mutations.

### Erg3 LoF is most likely not a *stand-alone* mechanism of echinocandin resistance

Variation in *ERG3* in the absence of acquired *FKS* mutations has been proposed as an independent driver of echinocandin resistance. Yet, this hypothesis is poorly supported by evidence. Scott *et al*. reported the emergence of an *ERG3*^*Q308K*^ mutation under micafungin monotherapy in serial clinical isolates of *C. lusitaniae*, although the majority of echinocandin-resistant isolates harbored *FKS1* mutations (18). Furthermore, both Rybak *et al*. (14) and later Hartuis *et al*. (19) reported that the dysfunction of Erg3, caused by a G111R mutation, led to an intermediate-to-resistant phenotype in *Candida parapsilosis*. Nevertheless, the *C. parapsilosis* species complex harbours a constitutive *FKS1*^*P660A*^ polymorphism that underlies reduced echinocandin susceptibility (16, 20). Rybak *et al*. suggested that *ERG3* LoF, combined with the *FKS1*^*P660A*^ mutation, results in the echinocandin-resistant phenotype (14). This hypothesis is supported by Papp *et al*. (15), who show how posaconazole exposure selects for an *ERG3*^*D14Y*^ mutation and azole and echinocandin cross resistance, in the ATCC 22019 background (15), which harbours the *FKS1*^*P660A*^ mutation (16). Additionally, Davari *et al*. (21) reported no mutations in *ERG3* or *FKS1*, apart from the naturally occurring *FKS1*^*P660A*^ mutation, in 105 *C. parapsilosis* isolates, of which only 3 (2.9%) were resistant to echinocandins. However, both resistant and susceptible isolates harboured the P660A substitution, supporting the notion that in *C. parapsilosis*, the *FKS1*^*P660A*^ polymorphism alone does not always lead to echinocandin resistance, but the combination of *FKS1*^*P660A*^ and *ERG3* variation does.

In *C. albicans*, the deletion of *ERG3* (*ERG3Δ*) in two different backgrounds (14), or partial deletion causing a frameshift in *ERG3* in a clinical isolate (22), did not result in echinocandin resistance. Similarly, a S258F mutation in *ERG3* in a clinical *C. tropicalis* strain was linked to azole and polyene but not echinocandin resistance (23).

To assess the role of Erg3 LoF in echinocandin resistance in *N. glabratus* and *C. auris*, we evaluated echinocandin susceptibility in wildtype (wt) and previously constructed *ERG3Δ* strains across three genetic backgrounds. **Figure 2** shows that the LoF of Erg3 does not confer resistance or decreased susceptibility to echinocandins in the studied backgrounds. In contrast, the Erg3 LoF in the *C. auris* Clade III background showed decreased tolerance to micafungin. Similarly, Carolus *et al*. (24) demonstrated that *ERG3*^*T308M*^ and *ERG3*^*G108\**^ mutations in *ERG11* LoF backgrounds were associated with collateral sensitivity to echinocandins, rather than resistance. Combined, these data suggest that the LoF of Erg3 alone does not lead to echinocandin resistance.

### Allosteric interactions between sterols and Fks might drive echinocandin resistance

The LoF of Erg3 drastically alters the membrane sterol composition, without necessarily imposing major fitness trade-offs (25). In *C. auris, ERG3Δ* strains have membranes enriched in ergosta-7,22-dienol, 4,14-dimethyl-zymosterol and lanosterol (24), whereas in *N. glabratus, ERG3Δ* results in the accumulation of ergosta-7,22-dienol, ergosta-5,7-dienol and fecosterol, with no detectable 4,14-dimethyl-zymosterol (25). Sterols are important for the integrity, fluidity, permeability, and function of the plasma and endomembranes and their associated proteins. In 2023, Hu *et al*. (2) characterized the structure and function of *S. cerevisiae* Fks, which is a large (200 kDa) transmembrane protein of approximately 1800 amino acids. Fks contains an extensive transmembrane (TM) domain with 17 helices (TM1-17), connected by several elongated loops. Hu *et al*. identified multiple orderly bound lipids in the TM domain, which are hypothesized to play an integral role in the conformational structure and function of the Fks protein. Interestingly, all three mutational hotspots (HS1, 2, and 3) are located on three neighbouring TM helices: TM5 (HS1), TM8 (HS2), and TM6 (HS3). The authors note that these HS regions are enriched in ordered lipids and that HS mutations, such as the S643P substitution in HS1, cause both conformational changes and lipid rearrangements (2).

Echinocandins are lipopeptides, and their lipid tail has been suggested to play an integral role in their inhibitory effect. Based on their findings, Hu *et al*. (2) proposed two possible echinocandin resistance mechanisms: either mutations may directly alter the echinocandin-binding site, involving interactions with the lipid tails of these lipopeptide drugs, or mutations may affect the response of Fks to membrane alterations induced by echinocandins, analogous to mechanisms described for other membrane-acting lipopeptide antibiotics like daptomycin and polymyxins (2).

Recently, an echinocandin-Fks binding model at atomic resolution was obtained by combining deep-mutational scanning of the three hotspots and Site Identification by Ligand Competitive Saturation (SILCS) – Molecular Dynamics (MD) simulations (1). The proposed model shows that several polar residues of Fks1 cause a deformation of the upper leaflet of the membrane and create a water-filled pocket between the three hotspots. As a result, parts of HS1 and HS2 are exposed to the extracellular solvent, which allows binding of the hydrophilic macrocycles of echinocandins. On the other hand, the lipophilic tails of anidulafungin and micafungin are embedded in the membrane and bind HS3, whereas the flexible tail of caspofungin, which is itself smaller, fits in a hydrophobic pocket between HS1 and HS2. The model supports the long-standing hypothesis that some substitutions most likely alter the shape of the binding site, thereby inhibiting drug binding, which ultimately leads to resistance. Importantly, this model highlights how echinocandin action takes place directly in and at the transmembrane interface. Therefore, it is reasonable to assume that binding and resistance are influenced by the specific lipid composition, organization, and fluidity of the cell membrane. Thus, we hypothesize that sterol composition changes, mediated by Erg3 LoF, could stabilize or modulate the interaction between Fks and echinocandins.

This sterol-protein interaction is likely highly dependent on the sterol composition. This idea is supported by the observation that other changes in sterol composition, for example, due to the LoF of *ERG6, NCP1*, and *ERG11*, lead to collateral sensitivity to echinocandins (25, 26), thus having an opposite effect to that of *ERG3* LoF. In addition, recently, Ross *et al*. (27) recently reported that the *FKS1*^*S639Y*^ mutation in *C. auris* confers collateral sensitivity to azoles, while the *FKS1*^*S639P*^, does not. The fact that a different substitution, at the same position, can impact the susceptibility to a drug that works by blocking ergosterol synthesis, again supports the hypothesis that the function of the essential Fks protein is potentially dependent on and influenced by membrane sterols. This also again stresses that epistasis of *ERG3* LoF and Fks variation is highly specific, depending on the *FKS* allele, accounting for (increased) resistance in some Fks variants, but having no effect or perhaps the opposite effect in other conformations, although the latter has not been reported.

Beyond the sterol-Fks interaction hypothesis, other mechanisms can be hypothesized or have been proposed. To investigate why *ERG3* mutations frequently emerged during experimental evolution under anidulafungin exposure, Ksiezopolska *et al*. (13) tested whether these mutations could affect stress tolerance or competitive fitness, but did not discern a clear effect. They also investigated whether *ERG3* mutations predated *FKS* mutations, or vice versa, by analyzing intermediate generations in their experimental evolution assay, finding no particular pattern, with mutations appearing first in one of the two genes, or simultaneously in consecutive generations. Although the outcomes of their experiments could not reveal a specific mechanism, they hypothesized that *ERG3* mutations might alter membrane composition, indirectly compensating for cell-wall changes caused by anidulafungin treatment.

Another hypothesis is that the LoF of Erg3 changes membrane lipid mobilisation or impacts lipid raft integrity, thereby altering the localization of Fks. Lipid rafts, rich in sterols and sphingolipids, are known to modulate the localization and activity of membrane-associated proteins, potentially affecting Fks functionality and their interaction with echinocandins.

Alternatively, sterol changes might impact the mobilisation of echinocandins directly. Recently, it was shown that caspofungin localizes to the vacuole where it is degraded, while anidulafungin concentrates at the cell surface, and rezafungin is partitioned between the surface and the vacuole (28).

Erg3 LoF could also influence the interaction between Fks and Rho1, a key regulatory GTPase required for β-1,3-glucan synthesis, which might indirectly modulate echinocandin susceptibility.

Finally, membrane sterol changes may alter broad cellular stress responses and signal transduction in pathways related to cell wall integrity or other functions.

All these hypotheses remain speculative and require further biochemical and biophysical investigations.

### Erg3 as a driver of multidrug resistance?

The suggested epistatic interplay between *ERG3* LoF mutations and *FKS* variation could have significant clinical implications. One of the most pressing potential consequences is the expansion of echinocandin resistance profiles, potentially broadening the spectrum of mutations that confer resistance beyond the traditional hotspot regions (HS1-3). The clinical case we describe illustrates this potential.

Additionally, numerous studies (8, 11-14, 18) have indicated that the combination of *ERG3* and *FKS* mutations frequently results in MDR or even pan-resistance, encompassing azole, polyene, and echinocandin resistance. Notably, *ERG3* LoF mutations often arise under echinocandin monotherapy (7, 8, 13), suggesting that *ERG3-FKS* mutagenesis may be a critical factor driving MDR evolution, even without prior exposure to multiple antifungal classes. This mechanism aligns with the known role of Erg3 in sterol biosynthesis, particularly its compensatory function in mitigating toxic sterol accumulation under azole pressure, and its influence on membrane sterol profiles that limit polyene efficacy (8, 24, 25).

It is important to not that beyond mutations, differential expression via transcriptional rewiring or aneuploidies, could also affect sterol-biosynthesis and thus echinocandin and multidrug resistance.

Despite the compelling evidence we provide here, the precise mechanism by which Erg3 LoF and Fks variation contribute to echinocandin resistance remains elusive. Future research should focus on validating these epistatic interactions at a molecular level, including investigations into how specific *FKS* variants interact with different membrane sterol compositions to confer resistance in some scenarios but collateral sensitivity in others. Ideally, systematic mutagenesis, dynamic molecular modelling and structural biology approaches should be combined to understand this complex mechanism, at a molecular and biophysical resolution. Understanding these epistatic interactions is critical, given that sterols and their biosynthesis underpin the modes of action of two-thirds of the primary antifungal drug class arsenal, while Fks is the target of the third major class.

## Methods

### Strains and growth media

The four *N. glabratus* strains depicted in **Figure 1** have been previously reported. The susceptible and resistant controls concern laboratory strain ATCC2001 and the clinical isolate from a cardiac valve infection in patient 14 reported by Diaz-Garcia *et al*. 2021 (29). The clinical isolates A and B are strains 7 and 8, respectively, reported by Diaz-Garcia *et al*. 2022 (3).

The *C. auris* and *N. glabratus* strains depicted in **Figure 3** have been previously reported too: the *C. auris ERG3*Δ strains were constructed by Carolus *et al*. 2024 (24) (the Clade I wt is strain B8441 (AR0387) and the Clade III wt is strain), and the *N. glabratus* wt strain (ATCC2001) and mutant (*ERG3*Δ-1) were constructed and reported before by Carolus *et al*. 2025 (25).

**Figure 3.**
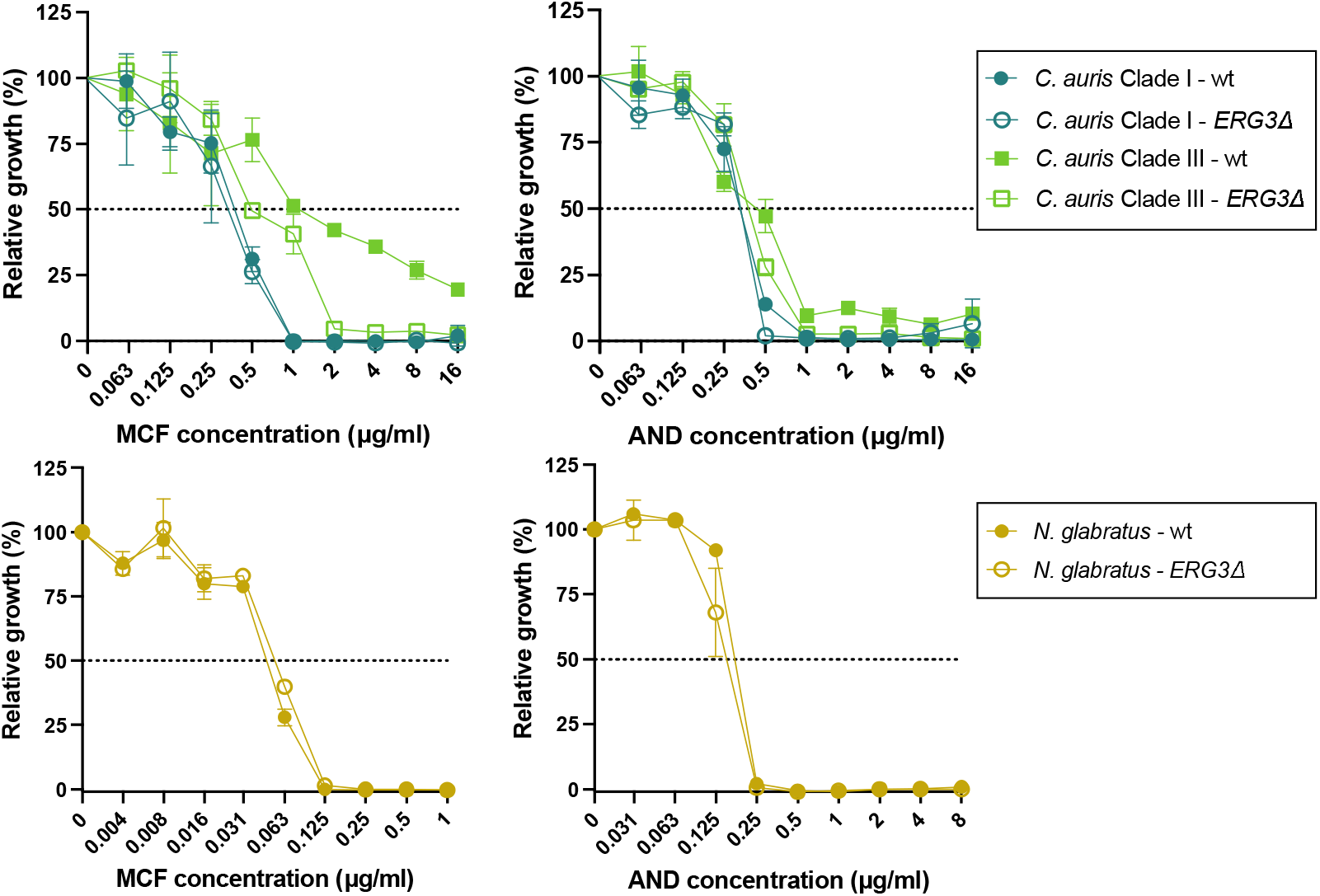
*In vitro* susceptibility testing of *ERG3*Δ strains in *C. auris* and *N. glabratus*. A broth dilution susceptibility test for micafungin (MCF) and anidulafungin (AND) was performed according to the EUCAST guidelines (4) on two independent cultures per strain.

All strains used in this study were stored at -80°C in 20% glycerol and routinely plated on solid YPD (1% yeast extract, 2% bacteriological peptone, 2% dextrose) agar (2%) at 37°C. Unless specified otherwise, cells were grown in MOPS (morpholinopropane sulfonic acid) buffered (pH 7) RPMI 1640 (Thermo Fisher Scientific) medium with 2% total glucose, at 37°C.

### Drug susceptibility testing

The EUCAST reference method (4) was used for AND and MCF broth dilution assays (BDA). Briefly, a twofold dilution range of the drug was prepared in a total volume of 200µL RPMI-MOPS (pH 7, 2% glucose, 1% DMSO) medium with approximately 20 000 cells (based on OD_600nm_ and serial dilution) in a round-bottom 96-well polystyrene microtiter plate (Greiner). Plates were incubated at 37°C for 24h (**Figure 1A**) and 48 hours (**Figure 3**), and growth was assessed spectrophotometrically (OD_600_). The growth cut-off of all MIC values from BDA was 50% growth compared to the drug-free control. Resistance breakpoints were determined based on EUCAST guidelines (5).

### *In vivo* colonization evaluation

8-week-old female BALB/c (Janvier) mice were immunosuppressed with dexamethasone (75 mg/kg IP) 3 days before and on the day of infection. An inoculum of 10^5^ cells in 100µL PBS was administered via tail vein injection. Treatment groups consisted of 5 mice per group. Each group was IP treated daily for 7 days, starting the day of infection (2h post inoculation), with one of two micafungin doses (1 mg/kg/day and 5 mg/kg/day) or the PBS vehicle (NTC; non-treated control). 8 days post infection, animals were sacrificed and colonization in kidneys, liver, spleen and brain was evaluated: organs were homogenized in 500µL sterile PBS with glass beads and shaking (20 seconds at 6m/second) in a FastPrep-24TM Classic lysis system (MP Biomedicals), homogenates were serially diluted and plated onto YPD agar for single colony (CFU) isolation after 48h incubation at 37°C.

### Whole-genome sequencing and data analysis

Genomic DNA from isolates A and B was extracted using the MasterPure yeast DNA purification kit (Lucigen) according to the manufacturer’s instructions. The purified genomic DNA was diluted to a concentration of 200 ng/μl in nuclease-free water, based on absorbance at 260 nm with a NanoDrop spectrophotometer (Isogen). Library preparation and sequencing were performed at Eurofins Genomics (Constance, Germany) on an Illumina NovaSeq6000 platform. Sequencing analysis (alignment, variant calling, and filtering) was performed as described in Carolus *et al*. 2025 (25). Briefly, quality control and trimming of the sequencing reads were performed using FastQC and Trimmomatic through the perSVade pipeline (version 1.02.6), which incorporates all tools used in the analysis (30). Trimmed reads were aligned to the *N. glabratus* ATCC2001 (CBS138) reference genome (version s02-m07-r35 from CGD) using BWA-MEM, and variant calling was performed using BCFtools, Freebayes, and GATK HaplotypeCaller (with ploidy parameter set to 1). Variants were retained only if supported by at least two of the three callers, with additional filtering based on read depth and allele frequency. High-confidence SNPs and indels were annotated using Ensembl Variant Effect Predictor (VEP), also implemented in perSVade.

To account for background genetic variation and determine likely recently acquired, isolate-specific variants, a maximum-likelihood phylogenetic tree was constructed using IQ-TREE based on SNPs detected in the clinical isolates. The isolates were placed within a previously defined phylogeny of 420 *N. glabratus* strains (31). An artificial background of clade-specific SNPs was constructed to filter out clade-associated variants from the clinical isolates. Only protein-altering variants present in less than 20% of clade members were retained for further analysis.

### ESM score calculation

To assess the potential functional impact of mutations, we computed ESM scores for orthologous sequences from five *Candida* species in **Table 1** using ESM variant (17) which leverages the ESM-1b protein language model (32). The variant effect score for each missense mutation is calculated as the difference in log-likelihood between the missense and the wild-type amino acid at the same position. For stop-gain variants, the effect score is defined as the lowest score among all possible missense mutations downstream of the stop codon within the lost protein region. A two-component Gaussian mixture model was fit to the distribution of ESM scores, excluding synonymous mutations. The threshold was set at the intersection point of the components to define very likely loss-of-function mutations. A histogram was generated using Python with the matplotlib and seaborn libraries.

## Data availability

The raw sequencing data of isolates A and B are available in the NCBI Sequence Read Archive under BioProject accession number PRJNA1258178. Retrievable via: https://dataview.ncbi.nlm.nih.gov/object/PRJNA1258178?reviewer=jhpp2f7eni8htlhli4q110diid

## Ethical statement

All animal experiments were approved in accordance with the ethical guidelines of the Ethics Committee of KU Leuven (project approval nr. 126/2022). Clinical strain information was obtained and published prior (3, 29) with Informed consent and ethical approval from the Hospital General Universitario Gregorio Marañón committee for medical ethics.

## Acknowledgements

This work was supported by the Fund for Scientific Research Flanders (FWO) under the framework of the JPIAMR – Joint Programming Initiative on Antimicrobial Resistance fund (project CycleDrug) granted to P.VD., and by a C3 grant from the Industrial Research Fund of KU Leuven (C3/22/007) granted to P.V.D. and K.L. H.C. was supported by a post-doctoral fellowships granted by KU Leuven Internal Funds (PDMT2/23/032) and the European Molecular Biology Organization - EMBO (ALTF 1105-2024). J.D. was supported by a predoctoral grant awarded by FIS (FI19/00021). J.D., P.E. and J.G. were supported by grants PI18/01155 and PI19/00074 from *Fondo de Investigación Sanitaria* (FIS. Instituto de Salud Carlos III; *Plan Nacional de I+D+I* 2017-2020) and by the European Regional Development Fund (FEDER) ‘A way of making Europe.’ V.B received funding from the European Union’s Horizon 2020 research and innovation program under the Marie Skłodowska-Curie grant agreement No 945352. S.J. and D.S., were supported by FWO PhD fellowships 11PRR24N and 11J8122N, respectively. The T.G group acknowledges support from the Spanish Ministry of Science and Innovation (grant numbers PID2021-126067NB-I00, CPP2021-008552, PCI2022-135066-2, and PDC2022-133266-I00), cofounded by ERDF “A way of making Europe”, as well as support from the Catalan Research Agency (AGAUR) (grant number SGR01551); “La Caixa” foundation (grant number LCF/PR/HR21/00737), and Instituto de Salud Carlos III (IMPACT grant IMP/00019 and CIBERINFEC CB21/13/00061-ISCIII-SGEFI/ERDF). A.P. and C.R.L. are supported by a Genome Québec and Genome Canada grant 6569. C.R.L holds the Canada Research Chair in Cellular Systems and Synthetic Biology. P.E. is recipient of a Miguel Servet contract supported by FIS (CPII20/00015) and J.G. is employed by Fundación para Investigación Sanitaria del Hospital Gregorio Marañón.

## Author contributions

H.C.: conceptualisation, investigation, formal analysis, visualisation, funding acquisition, supervision and writing original draft. J.D., V.B., S.J., D.S., C.L.R., L.V., and A.P.: investigation. C.R.L., K.L., T.G., P.E., J.G. and P.V.D.: supervision and funding acquisition. All authors edited and/or approved the manuscript.

## Supplementary

**Table S1.**
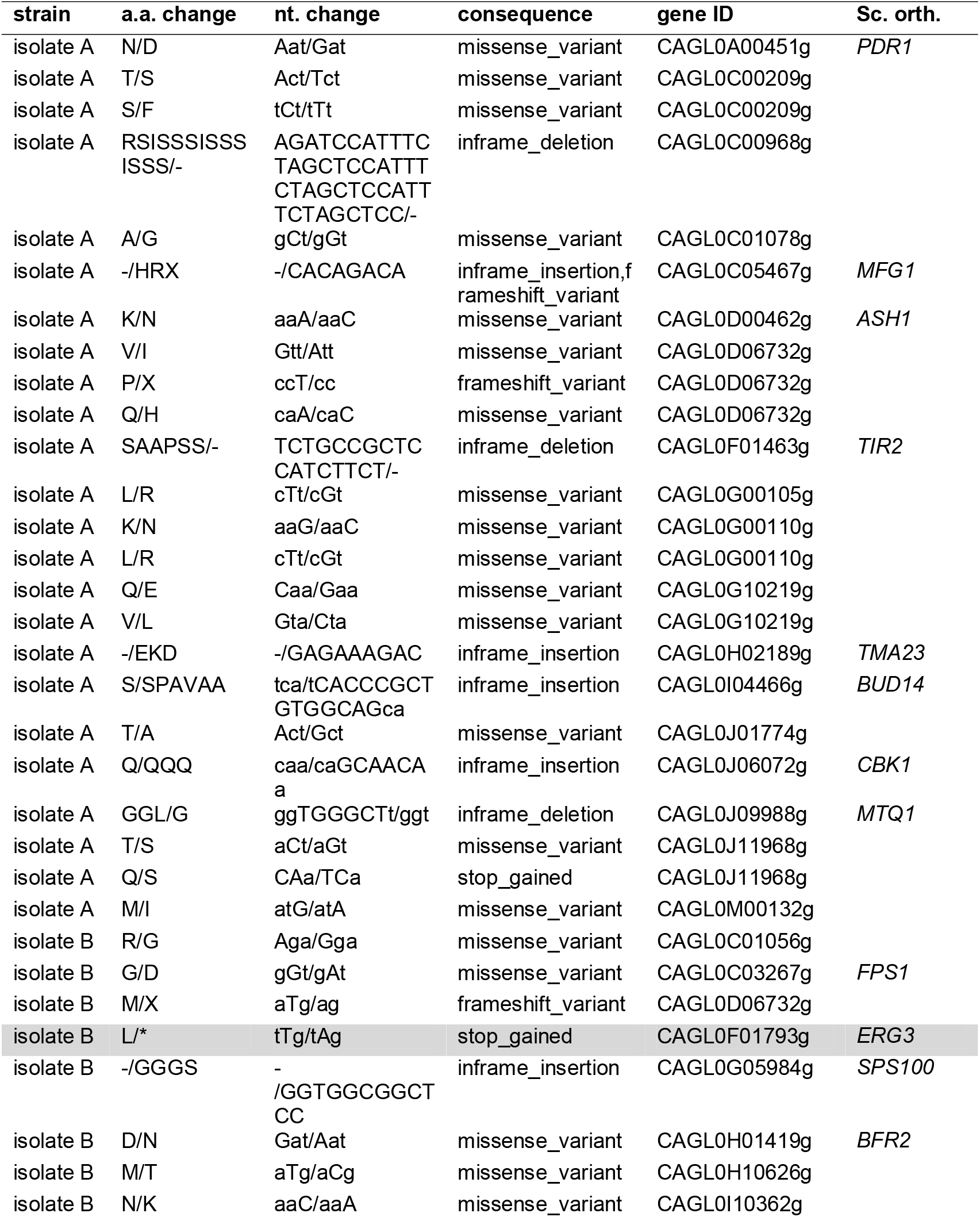
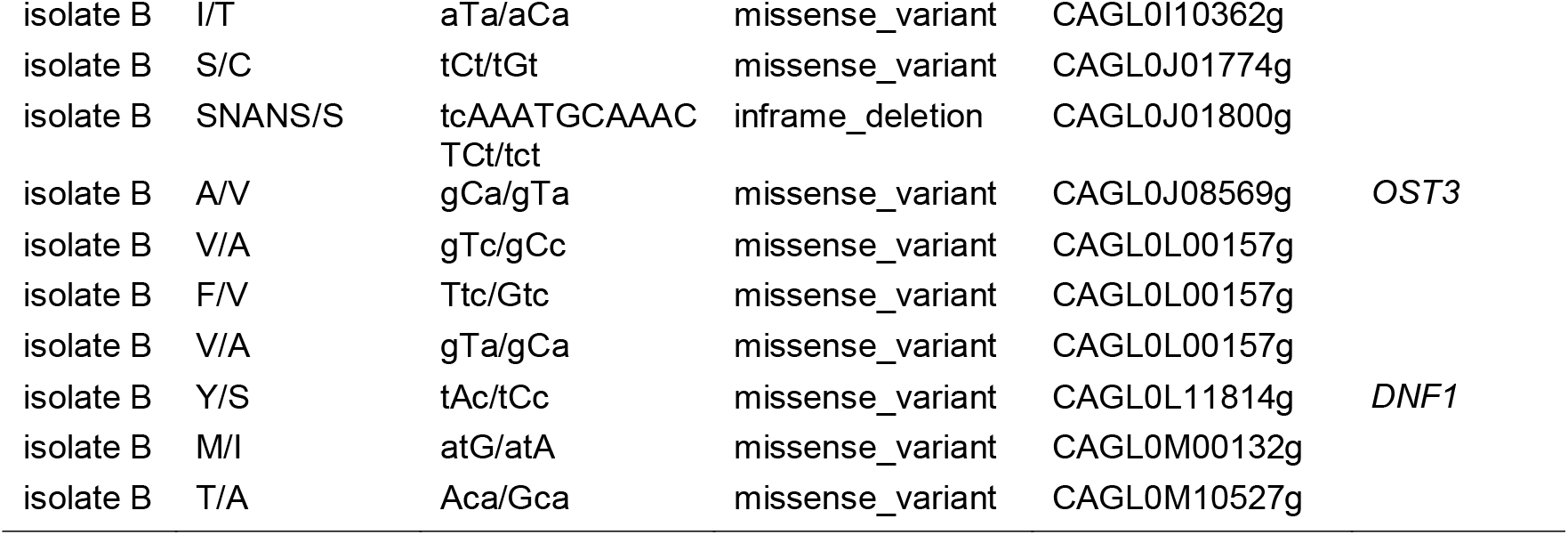
Unique variants identified between isolate A and isolate B in whole-genome sequencing analysis. For each variant, the amino acid change (a.a. change), nucleotide change (nt. change), consequence (type of mutation), gene ID in the reference genome used in this study, and corresponding *S. cerevisiae* ortholog name (Sc. orth.) is given. The mutation in *ERG3* is highlighted in grey.

